# Relationship between epicardial and perivascular fatty tissue and adipokine-cytokine level in coronary artery disease patients

**DOI:** 10.1101/470872

**Authors:** Olga Gruzdeva, Evgenya Uchasova, Yulia Dyleva, Daria Borodkina, Olga Akbasheva, Viktoria Karetnikova, Natalia Brel, Kokov Alexander, Olga Barbarash

**Author notes:** **Corresponding author:** Evgenya Uchasova, Federal State Budgetary Institution, Research Institute for Complex Issues of Cardiovascular Disease, Laboratory of Research Homeostasis, 6 Sosnovy Bvld, Kemerovo 650002, Russian Federation.

## Abstract

The aim of this study was to determine the relationship between the thickness of EAT and PVAT and the adipokine-cytokine profile of patients with coronary heart disease, which can be of significant importance for predicting the course of CVD. 84 patients with CVD, were assessed and divided into two groups based on the presence of visceral obesity (VO). In VO patients, the thickness of the epicardial deposits of the left and right ventricles were 1.75 and 1.43 times greater, respectively, than in patients without VO. For patients with VO, the prevalence of the volume of the left anterior descending artery was 10% higher, and the middle third of the envelope artery was 28% higher, when compared to patients without VO. When evaluating inflammatory status, it was established that the concentration of TNF-α and IL-1β, leptin in the blood serum of patients with VO exceeded the values of patients without VO. Level of proinflammatory IL-10 was 2-times lower in patients with VO. The findings of this study show that the increase of EAT and PVAT are independent risk factors of CVD, as well as a possible model for the assessment of drug effectiveness for CVD.

## Introduction

Obesity is a rapidly growing problem that is becoming an epidemic on a global scale, affecting both children and adults [1,2]. This condition is defined as the result of the formation of abnormal or excessive fatty deposits, which can be harmful to a person’s health [3]. An important factor in understanding the cardiovascular effect of an increased amount of adipose tissue (AT) has been the recognition that adipocytes are endocrine and paracrine determinants of vascular function. With obesity, AT can become dysfunctional, which is accompanied by ectopic fat deposits in other tissues that regulate metabolic homeostasis [4]. Expansion of AT is associated with numerous “local” consequences, including inflammation [5], fibrosis [6], hypoxia [7], dysfunctional adipokine secretion [8], and impaired mitochondrial function [9]. At the biochemical level, AT dysfunction includes abnormal metabolism of glucose and lipids, insulin resistance, activation of the renin-angiotensin system, hypercoagulation, inflammation, and endothelial dysfunction, which provide important mechanisms linking obesity to cardiovascular disease (CVD).

The wide introduction of diagnostic imaging methods allows for the classification of obesity according to visceral and subcutaneous tissues, depending on the location of excessive fatty deposits [10]. Many studies have demonstrated that visceral obesity (VO) is associated with an increased risk of morbidity and mortality from CVD, including stroke, congestive heart failure, and myocardial infarction (MI) [11,12]. Thus, despite the similarity in terms of structure, the regional location of fat deposits has a significant impact on the function and properties of the heart. Accumulation of fat in the visceral region is not the only metabolically active deposit, as at least 6 more regional deposits are characterized by similar disorders in the background of chronic inflammation. There is increasing evidence regarding the effect of epicardial AT (EAT) and perivascular AT (PVAT) localization on the risk of developing CVD [13,14].

The aim of this study was to determine the relationship between the thickness of EAT and PVAT and the adipokine-cytokine profiles of patients with coronary artery disease (CAD), which can be of significant importance for predicting the course of CVD.

## Materials and methods

### Ethical considerations

The study protocol was approved by the Local Ethics Committee of the Federal State Budgetary Institution Research Institute for Complex Issues of Cardiovascular Diseases, and was developed according to World Medical Association’s Declaration of Helsinki on Ethical Principles for Medical Research Involving Human Subjects, 2000 edition, and the “GCP Principles in the Russian Federation”, approved by the Russian Ministry of Health. Patients were included in the study after they provided written informed consent.

### Study subjects

Eighty-four patients with coronary artery disease (CAD), comprising of 65 men and 19 women, whose mean age was 58.7 years (range, 52.2;63.5 years), were included in this study. Diagnosis of CAD was established according to the criteria of the All-Russian Scientific Society of Cardiology (2007) and the European Cardiology Society (2013), American College of Cardiology Foundation, American Heart Association, and the World Heart Federation Task Force for the Redefinition of Myocardial Infarction [15]. Patients included in the study received standard antianginal- and antiplatelet-agent therapy. Patients were excluded due to the following criteria: patients younger than 50 and over 80 years-old; having a history of diabetes mellitus; level of glycosylated hemoglobin (HbA1c) of >6.0%; and the presence of severe concomitant pathologies, including oncological, infectious, mental disease, chronic obstructive pulmonary disease, connective tissue disease, and renal and/or hepatocellular insufficiency.

### Imaging assessments

In patients, the presence of visceral obesity (VO) was verified by measuring the area of PVAT and subcutaneous adipose tissue by means of multi-spiral computed tomography (MSCT) on a Siemens Somatom 64 computed tomography scanner (Siemens Healthcare, Erlangen, Germany). For VO, the area of PVAT should be greater than 130 cm^2^, and the ratio of the area of the visceral and subcutaneous adipose tissue (VAT / SAT) should be greater than or equal to 0.4 [16]. The determination of the thickness of the EAT is performed by magnetic resonance imaging (MRI) on an Exelart Atlas 1.5-T MR imager (Toshiba, Tokyo, Japan). Thickness measurements of EAT was carried out on images oriented along the short axis of the heart. EAT thickness was measured at 3 points along the anterior wall of the right ventricle, with the average value then calculated (Figure 1). In addition, the same method was used to measure EAT thickness at the back wall of the left ventricle, followed by the calculation of the mean value (Figure 2). Determination of the volume and thickness of the SCAT at different anatomical locations was performed using the MSCT method with the following parameters: 1-mm axial slice thickness, an image matrix of 512×512, tube voltage of 120 kV, and a tube current of 100 mA. The MSCT images were acquired using bolus contrast, followed by a quantitative evaluation.

**Figure 1.**
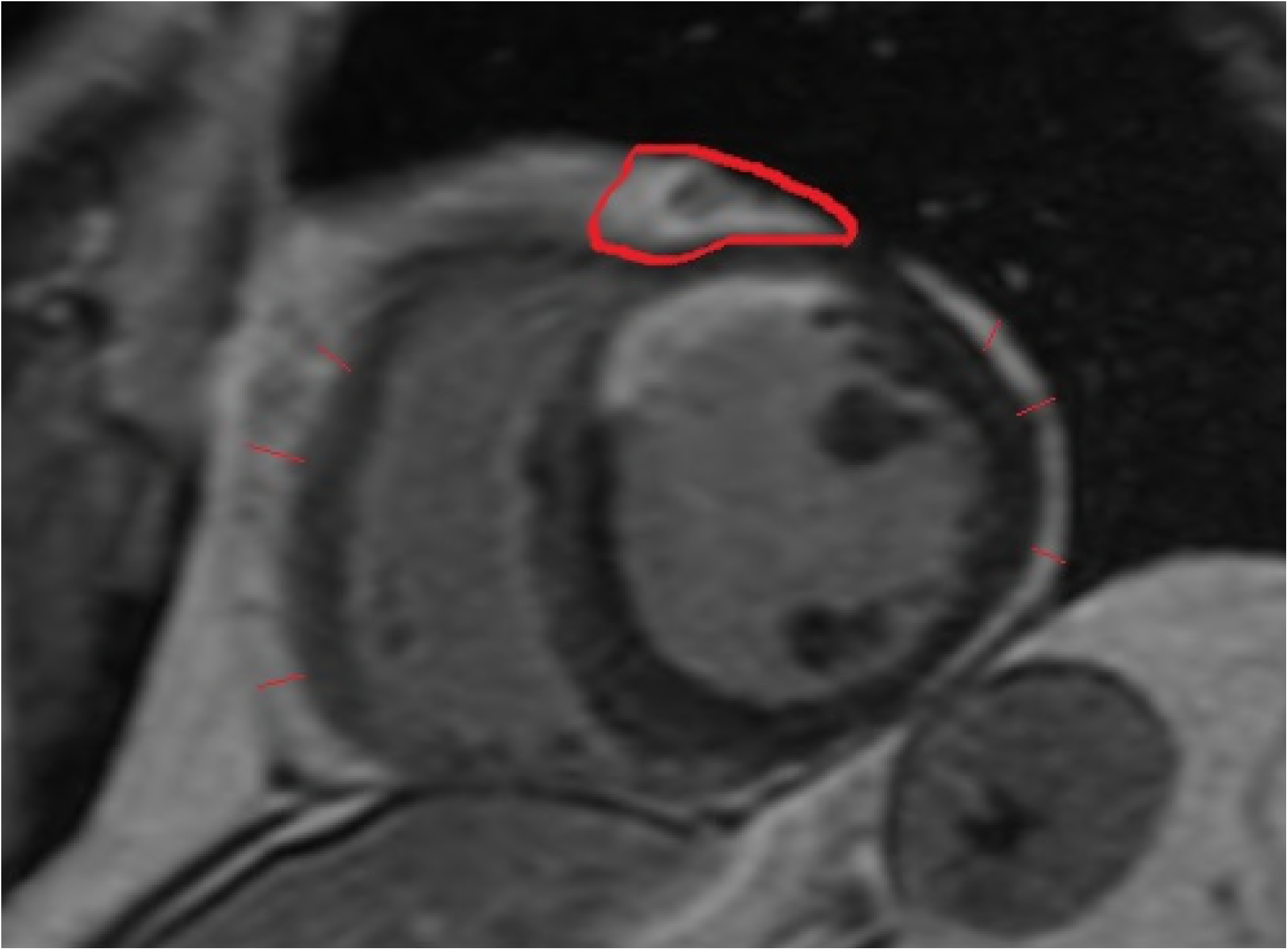
Quantitative assessment of the thickness of epicardial adipose tissue along the anterior wall of the right ventricle, as well as the thickness of the epicardial adipose tissue along the posterior wall of the left ventricle. Measurement of the area of the epicardial adipose tissue at the level of the coronal sulcus (red contour).

**Figure 2.**
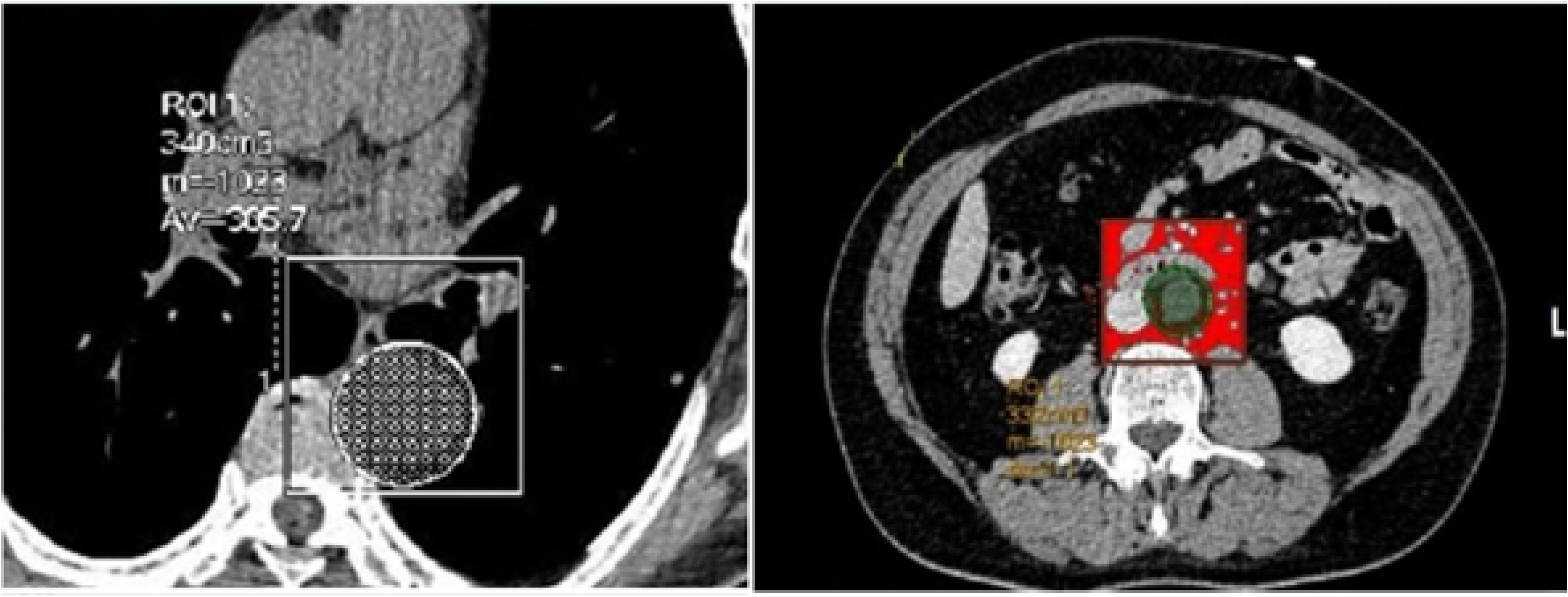
Quantitative assessment of the volume of para-aortic adipose tissue at the level of the thoracic and abdominal aorta (axial image).

Determination of the volume of the para-aortic adipose tissue at the level of the thoracic aorta was performed at the level of the bifurcation of the pulmonary artery by 70±1 mm in the caudal direction; the volume of the abdominal aorta was determined at the level of the L2–L5 vertebrae over 70±1 mm from the abdominal aortic bifurcation in the cranial direction. In addition, the thickness of the paracoronal vessels was assessed on post-contrast imaging at the level of the left coronary artery and at the level of the proximal and middle segments of the anterior descending envelope and right coronary arteries (Figure 3).

**Figure 3.**
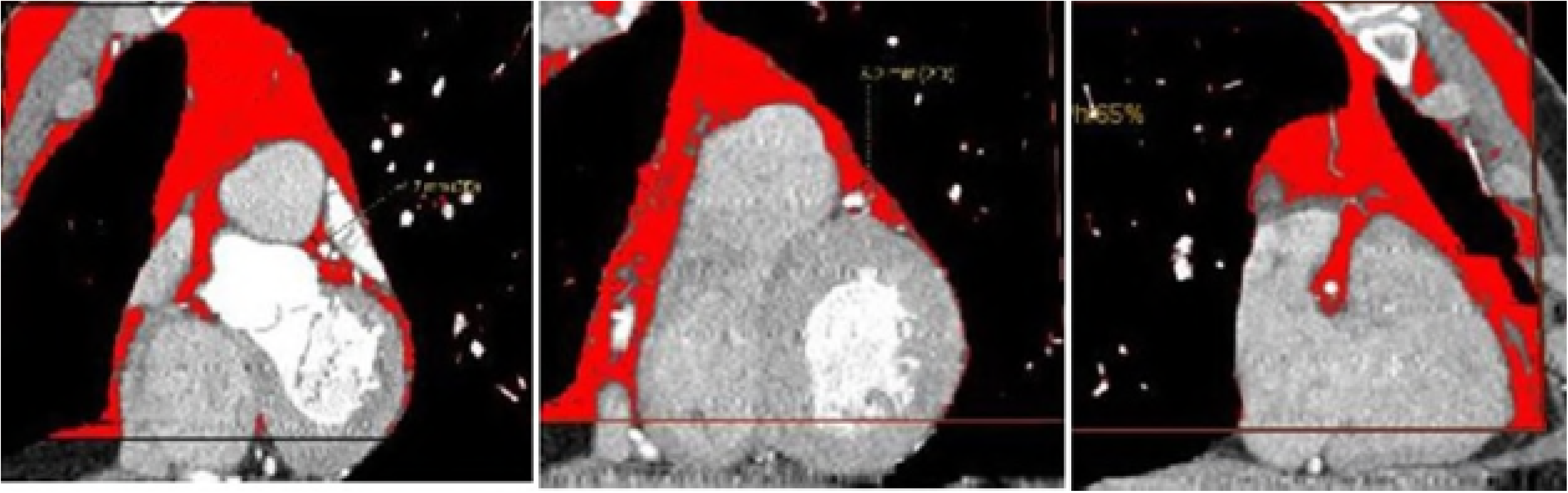
Quantitative assessment of paracoronary fat issue at the level of the proximal segment of the anterior descending artery, the middle third of the anterior descending artery, and the proximal segment of the right coronary artery.

### Serum sample evaluations

The serum total cholesterol (TC), triglycerides (TGs), very low-density lipoprotein cholesterol (VLDL-C), low-density lipoprotein cholesterol (LDL-C), high-density lipoprotein cholesterol (HDL-C), apolipoprotein (Apo)-A1, Apo-B, free fatty acid (FFA), glucose, and glycated hemoglobin (HbA1c) levels were measured using enzymatic methods with end points on an automatic biochemical analyzer (Konelab 30i, Thermo Fisher Scientific, Waltham, MA). The serum insulin and C-peptide levels were measured using commercially available ELISA kits (Monobind Inc., Lake Forest, CA). Insulin sensitivity was evaluated based on insulin and fasting glucose levels using the homeostasis model assessment (HOMA) index as follows: insulin level × glucose level/22.5. Insulin resistance (IR) was diagnosed when the HOMA-IR index was >2.77 [14].

The leptin, soluble receptor (sOB-R), and adiponectin levels, as well as the serum sample levels, were measured using commercially available enzyme immunoassay kits (BioVendor, Brno, Czech Republic; eBioscience, Vienna, Austria). Each sample was tested twice and intra-assay correlations of variation were calculated. Samples averaged 5.8% with all standard curve correlation coefficients >0.998. The levels of interleukin (IL)-1β, IL-10, and tumor necrosis factor-α (TNF-α) were measured using a commercially available enzyme-linked immunosorbent assay (ELISA) (Bender MedSystems GmbH, Vienna, Austria) (coefficient of variation [CV], 7.03–8.99%). The free leptin index (FLI) is defined as the ratio of the leptin to sOB-R; hence, the sOB-R level provides an indication of the free leptin level. Leptin resistance was diagnosed when the serum FLI was >0.25.

### Statistical analyses

The statistical analyses were performed using Statistica software, version 6.1 (Dell Software Inc., Round Rock, TX). The Mann-Whitney U test was used to compare the non-normally distributed independent variables. Spearman’s rank correlation analysis was used to identify the dependencies between the variables. The data are presented as the median values and the 25th and 75th quartiles. A value of *p* < 0.05 was considered statistically significant.

### Results

The clinical and demographic data of the study patients are shown in Table 1. Patients were divided into 2 groups, based on whether they had VO present; the first group included 54 patients with VO, the second group included 30 people without VO (Table 2). Patients were comparable in age, the presence of risk factors for CVD (i.e., hypertension, smoking, angina pectoris, dyslipoproteinemia, and congestive heart failure), and myocardial infarction in the anamnesis.

According to the MRIs and MSCTs of patients with VO, the accumulation of fat in epicardial and paravascular adipocytes was more pronounced, compared with patients without VO. In VO patients, the thickness of the epicardial deposits of the left and right ventricles were 1.75 and 1.43 times greater, respectively, than in patients without VO (Figure 4). A similar pattern was observed in the visualization of the intrauterine tract of the abdominal aorta, such that the volume of the fat deposits in patients with VO was 1.3 times greater than the volume of the PVAT in patients without VO. However, for the thoracic compartment there was no difference in the volume of periortal adipose tissue between the 2 patient groups. For patients with VO, the prevalence of the volume of the left anterior descending artery was 10% higher, and the middle third of the envelope artery was 28% higher, when compared to patients without VO. Whereas in patients without VO, the prevalence of fatty deposits around the right coronary artery and the lower third of the envelope artery was 6.7% and 11.4% higher, respectively, when compared to patients with VO.

**Figure 4.**
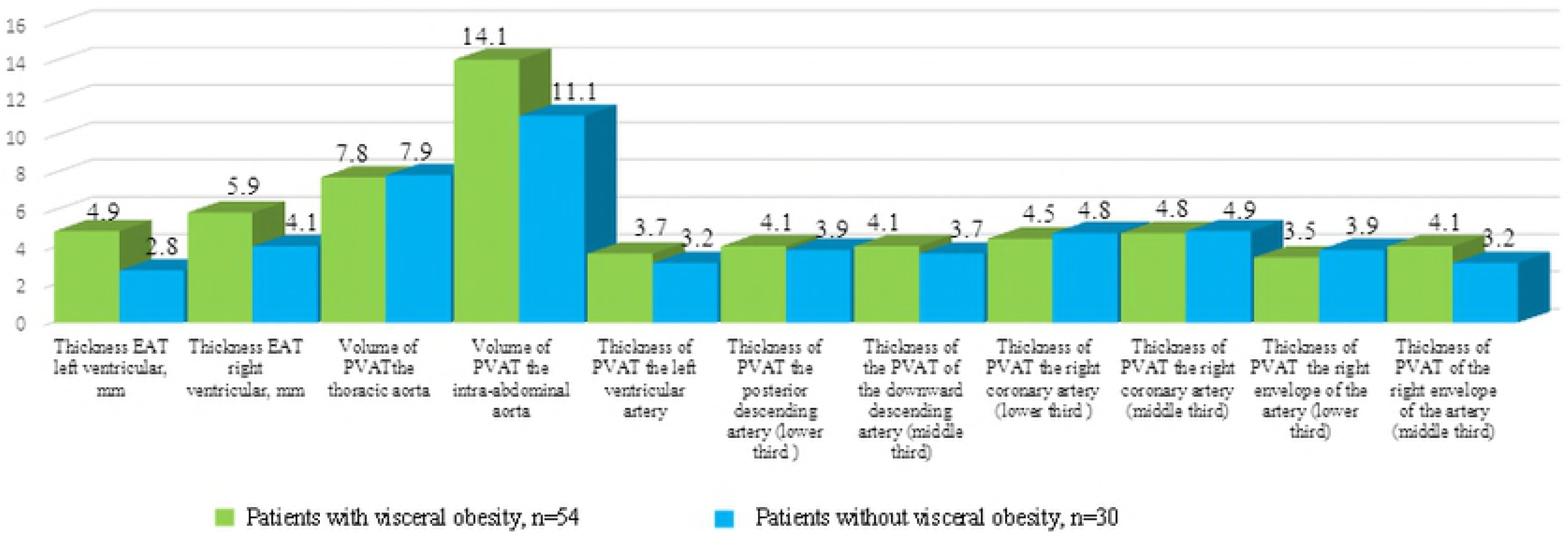
Quantitative evaluation of EHT and HPLC in the study groups.

In the group of patients with VO, a direct correlation was found between the area of PVAT and the thickness of the epicardial deposits of the left ventricle (r = 0.26, *p* = 0.02), thickness of the epicardial deposits of the right ventricle (r = 0.26, *p* = 0.01), and the volume of both the chest PVAT (r = 0.51, *p* = 0.0001) and the abdominal aorta (r = 0.62, *p* = 0.0002). The volume of fat deposits of the paracoronary arteries did not depend on the magnitude of the PVAT or the volume of the PVAT of the aorta. However, there was a direct connection between both PVAT of the left ventral descending artery and epicardial deposits of the left ventricle (r = 0.64, *p* = 0.00), as well as PVAT of the right ventricular artery with epicardial deposits of the right ventricle (r = 0.26, *p* = 0.02).

It was discovered that the concentration of atherogenic lipid metabolism values (LDL-C, LD-VLDL, Apo-B, TAG, Apo-B/Apo-A1, FFA) in the blood serum of patients with CAD on the background of VO patients increased significantly and the content of anti-atherogenic fractions decreased (HDL-C and Apo-A1), relative to patients without VO (Table 3). In patients with CAD, changes in carbohydrate metabolism were more pronounced in patients with VO; in particular, an increase in the HOMA index by a factor of 1.5 and the presence of IR, compared with patients without VO (Table 4). In addition, patients with VO are characterized by hyperinsulinemia, an increase in the concentration of C-peptides, compared to patients without VO. The concentration of glucose and glycated hemoglobin did not differ significantly (*p* > 0.05).

The presence of atherogenic dyslipidemia and IR was accompanied by the formation of widespread atherosclerotic vascular lesions in patients with CAD and VO. Based on the results of angiographic examinations and the duplex scanning of blood vessels, isolated CAD was recorded in 18.5% (n = 10) of patients with VO and 33.3% (n = 10) of patients without VO (*p* = 0.04). The defeat of several basins with stenoses of less than 30% was found in 44.4% (n = 24) of patients with VO and in 30.0% (n = 10) of patients without VO (*p* = 0.04). The lesions of several basins with stenoses from 30–50% was detected in 27.8% (n = 15) of patients with VO and in 16.7% (n = 5) of patients without VO (*p* = 0.04). A lesion of several basins with stenoses of more than 50% was recorded in 9.3% (n = 5) of patients with VO and in 13.3% (n = 4) of patients without VO (*p* = 0.26). In addition, 3 or more lesions of the coronary vessels predominated, which occurred in 66.7% (n = 36) of patients with VO, compared to 46.7% (n = 14) in patients without VO (*p* = 0.03).

As a result of the evaluation of the adipokine balance parameters in the blood serum of patients with CAD (Table 5), it was found that for patients with VO, the level of leptin was 1.67 times higher, and the concentration of sOB-R was 32.5 % lower, compared to patients without VO. Leptin resistance was confirmed by FLI, which was 2.5 times higher in patients with VO, compared to patients without VO. On the contrary, the concentration of adiponectin in the serum of patients with VO was 27.7% lower than that of patients without VO. When evaluating inflammatory status, it was established that the concentration of TNF-α and IL-1β in the blood serum of patients with VO exceeded the values of patients without VO by 1.5 and 2.0 times, respectively. Additionally, the level of proinflammatory IL-10 was 2.2-times lower in patients with VO, when compared to patients without VO.

The correlation analysis confirmed the relationship between EAT thickness and the serum concentrations of adipokines. Thus, in patients with VO, the leptin concentration had a negative dependence on the thickness of the left and right ventricular (r = −0.28, *p* = 0.02; r = −0.33, *p* = 0.02, respectively). A negative dependence on the EAT thickness was also established for the FLI (r = −0.28; *p* = 0.03). In addition, the PVAT area was directly dependent on the leptin concentration (patients with VO, r = 0.48, *p* = 0.02; patients without VO, r = 0.33, *p* = 0.02). A similar dependence was established for FLI (patients with VO, r = 0.28, *p* = 0.03; patients without VO, r = 0.22, *p* = 0.04).

Adiponectin levels were inversely related to the PVAT value (patients with VO, r = −0.43, *p* = 0.00; patients without VO, r = −0.18, *p* = 0.03). A general finding for both patients with and without VO was a positive correlation between the concentration of TNF-α, as well as IL-1β, and the area of VAT (patients with VO, r = 0.24, *p* = 0.05; r = 0.46, *p* = 0.04, respectively; patients without VO, r = 0.33, *p* = 0.01; r = 0.34, *p* = 0.02, respectively). The obtained results indicate the absence of the identity of the metabolic processes in EAT and PVAT, despite the direct correlation with EAT thickness and the number of PVAT.

It should be noted that the PVAT of the abdominal aorta is similar to the properties of visceral AT. An analysis of the possible relationship between paravascular adipocytes and adipokine exchange rates showed a direct relationship between the volume of the abdominal aortic ventricle and leptin (r = 0.44, *p* = 0.01), as well as the FLI (r = 0.56; *p* = 0.03). In addition, a link was also found with lipid metabolism indices, namely, FFA (r = 0.48, *p* = 0.02). A direct relationship between the adipose tissue thickness of the left coronary artery trunk and TNF-α (r = 0.88, *p* = 0.01), the circumflex artery (r = 0.88, *p* = 0.01), and glucose levels (r = 0.49; *p* = 0.03). In patients without VO, such connections were not established.

Using logistic regression analysis, it was established that of all the variables studied, the most influential variables on EAT thickness were VO (OR = 1.9, 95% CI [1.6–2.8]), LV hypertrophy (OR = 1.3; 95% CI [1.1–1.8]), HOMA index (OR = 0.8, 95% CI [0.6–1.1]), and FFA (OR = 1.2, 95% CI [1.1–1.8]). The most significant factors for increasing the volume of the coronary artery were TNF-α (OR = 1.5, 95% CI [1.1–1.9]), adiponectin (OR = 0.9, 95% CI [0.6– 1.1]), and leptin (OR = 1.2, 95% CI [1.1–1.3]). Over the course of the analysis, there was no evidence of a correlation between VAT magnitude and the thickness PVAT of the coronary artery.

### Discussion

This study examined the relationship between both EAT and PVAT, and the adipokine-cytokine profile of patients with CAD. According to the literature, EAT is a type of visceral adipose tissue localized near the myocardium and around the coronary arteries [17]. PVAT and EAT have the same embryological origin, and the increase in the size of either fat store is associated with the calcification of the coronary arteries [18,19], as well as the development of CAD [20]. Some researchers believe that an increase in the thickness of epicardial fat reflects the presence of visceral obesity in the body and serves as a prognostic marker of coronary heart disease and its associated complications [21]. The results obtained in this study show that EAT thickness is directly dependent on the value of the PVAT; this concurs with the results of previous studies [22,23]. Our findings also show that the presence of a direct connection between an increase in EAT thickness and LV hypertrophy, as well as insulin resistance. Similar properties are also present with PVAT [18,24].

Despite the presence of this bond, epicardial adipocytes have unique properties that distinguish them from the fat cells of other depots. EAT in healthy individuals is represented mainly by brown AT, whereas PVAT is white [19]. The cells of white and brown AT differ significantly from each other: the cell of white adipose tissue has inside itself one large fatty vial. It occupies almost the entire cell and pushes its core to the periphery, which becomes flattened. In adipocyte brown AT, there are several small fat drops and many mitochondria that contain iron (in cytochromes), which is responsible for the brown color of the tissue. Under physiological conditions, adipocytes of EAT perform a number of important functions for the myocardium: metabolic (absorb excess FFA and act as a source of energy under ischemic conditions), thermogenic (protect the myocardium from overheating), and mechanical, as well as the synthesis of adiponectin and adrenomedullin-possessing cardioprotective properties [20,25].

However, against the background of obesity and the progression of coronary atherosclerosis, the phenotype of EAT adipocytes from brown to white changes due to activation of the IL-6 signaling pathway of JAK-STAT3 [26]. For white PVAT adipocytes, obesity is characterized by intense lipolysis with the formation of FFA, as well as an increase in the production of pro-inflammatory factors (IL-1β, IL-6, TNF-α) and leptin, which enters the bloodstream and causes irreversible changes in the body (i.e., dyslipidemia and insulin resistance). Thus, for patients with CAD, the presence of IR and hyperinsulinemia in patients with VO, as well as atherogenic dyslipidemia with an increase in the concentration of FFA, TG, and VLDL, were characteristic of these patients when compared to those without VO. In addition, for patients with coronary artery disease and VO, against hypertension, hyperleptinemia, leptin resistance and reduction in adiponectin concentration were recorded.

However, the metabolic processes accompanying hypertrophy and changes in the phenotypes of EAT adipocytes differ from those in visceral adipocytes. Therefore, we have demonstrated that an increase in PVAT is associated with leptin hypoproducts against adiposopathy, as well as with the development of leptin resistance. While the EAT thickness was inversely related to the concentration of leptin and the FLI index (one of the main leptin-resistance markers).

Perivascular fat is located around vessels of different sizes, does not have barriers separating it from the adventitia of the vessel [26], therefore, this results in synthesized cytokines and chemokines can act directly on the vascular wall, potentiating vasospasm, endothelial dysfunction, proliferation of smooth muscle cells, migration of leukocytes in the intima, and fibrosis [27]. A large amount of data supports regional, phenotypic, and functional differences between deposits of different localizations [28]. The results obtained in this report confirm this assumption. We did not find a correlation between PVAT thickness and the thickness of the coronary arteries; while for EAT thickness, this relationship was recorded and was of a direct consequence. Interestingly, the PVAT coronary artery is slightly anomalous, because it resembles the phenotype of white AT, but includes adipocytes of different sizes and differentiated statuses, similar to brown AT [29].

EAT and PVAT coronary arteries have different phenotypes, but the presence of a common microcirculation network and direct anatomical proximity allows these tissues to interact and influence each other. Therefore, the thickness of the perivascular deposits of the left coronary artery depended only on EAT thickness (right ventricle, which it directly contacts with), and the vessels of the right side of the heart from the EAT (left ventricle). Evaluation of the volume of paraortal AT showed no difference in the thoracic region, depending on the presence of VO. Perhaps this is due to the thoracic aorta being surrounded by brown adipocytes, of which the number and volume do not change over a patient’s life or the development of obesity. While the volume of the abdominal region, as represented by white AT, was directly dependent on PVAT and was greater in patients with VO.

The heterogeneity of the PVAT of a vessel can be important for the realization of its protective or pro-atherogenic function. In the course of our study, it was shown that the metabolic potential of the abdominal aorta is similar to that of the thoracic aorta, as evidenced by the correlations with the concentration of FFA, leptin, and FLI, while for the thoracic region such connections were not discovered. In addition to the presence of traditional correlations for visceral deposits with TNF-α and leptin, the thickness of the left ventricular artery stenosis depended on the concentration of adiponectin. Previously, such connections were demonstrated only in subcutaneous adipose tissue, whereas for other visceral deposits such connections are not characteristic.

## Conclusion

The findings of this study show that the increase of EAT and PVAT are independent risk factors of CVD, as well as a possible model for the assessment of drug effectiveness for CVD. The study of the molecular basis of PVAT and EAT function can provide a more complete understanding of the etiopathogenetic mechanisms of CVD and develop an effective strategy for their prevention and control.

## Supporting information

**S1 Table 1. Clinical and anamnestic characteristics of the study patients**.

**S2 Table 2. Baseline clinical characteristics of the study patients, based on the presence of visceral obesity**

**S3 Table 3. Parameters of lipid metabolism in patients with coronary artery disease, depending on the presence of visceral obesity**

**S4 Table 4. Parameters of carbohydrate metabolism of the study patients, depending on the presence of visceral obesity (Me: 25;75)**.

**S5 Table 5. Level of blood serum adipokines and cytokines from coronary artery disease patients, depending on the presence of visceral obesity (Me: 25; 75)**.

**S6 Figure 1. Quantitative assessment of the thickness of epicardial adipose tissue along the anterior wall of the right ventricle, as well as the thickness of the epicardial adipose tissue along the posterior wall of the left ventricle. Measurement of the area of the epicardial adipose tissue at the level of the coronal sulcus (red contour)**.

**S7 Figure 2. Quantitative assessment of the volume of para-aortic adipose tissue at the level of the thoracic and abdominal aorta (axial image)**.

**S8 Figure 3. Quantitative assessment of paracoronary fat tissue at the level of the proximal segment of the anterior descending artery, the middle third of the anterior descending artery, and the proximal segment of the right coronary artery**.

**S9 Figure 4. Quantitative evaluation of EAT and PVAT in the study groups**

